# Seasonal variation of behavioural thermoregulation in the Spotted Salamander (*Ambystoma maculatum*)

**DOI:** 10.1101/2024.03.15.585242

**Authors:** Danilo Giacometti, Glenn J. Tattersall

## Abstract

Mounting evidence suggests that temperature seasonality plays a pivotal role in shaping the thermal biology of ectotherms. However, we still have a limited understanding of how amphibians maintain thermal balance in the face of varying temperatures, especially in fossorial species. Due to thermal buffering underground, theory predicts relaxed selection pressure over thermoregulation in fossorial ectotherms. As a result, fossorial ectotherms typically show low thermoregulatory precision and low evidence of thermotactic behaviours when tested in laboratory thermal gradients. In this study, we evaluated how temperature selection (*T*_sel_) and behavioural thermoregulation differed between seasons in the fossorial Spotted Salamander (*Ambystoma maculatum*). By comparing thermoregulatory parameters between the activity and overwintering seasons, we provide evidence that *A*. *maculatum* engages in active behavioural thermoregulation despite its fossorial habit. In both seasons, we found *T*_sel_ to be consistently offset higher than prevailing thermal conditions. Thermoregulation differed between seasons, with salamanders having higher *T*_sel_ and showing greater evidence of thermophilic behaviours in the active season compared to the overwintering season. Our study highlights that the combination of behavioural and thermal biology measurements is a necessary step to better understand the mechanisms that underlie body temperature control in amphibians. Ultimately, our study provides a broader understanding of thermoregulation in amphibians, particularly in the context of behavioural responses to seasonality in fossorial species.

**Summary statement:** By comparing thermoregulatory parameters between seasons, we demonstrate that the Spotted Salamander engages in active behavioural thermoregulation despite being fossorial.

## Introduction

Ectotherms depend on external sources of heat to regulate body temperature (*T*_b_) within favourable values because of their relatively low ability to generate and retain endogenous heat (Angilletta, 2009). Shifts in environmental temperatures affect organismal processes of ectotherms at all levels of organisation, from energy turnover (Whitford and Hutchison, 1963) to locomotion (Greenberg and Palen, 2021). To maintain thermal balance, an ectotherm must be able to sense and process the range and amplitude of environmental temperature variation. Coordinated physiological and behavioural processes are then put in action to modulate heat transfers between the organism and its surroundings (Tattersall et al., 2012). In heliothermic lizards, for instance, an increase in heart rate and peripheral blood flow have been shown to improve the delivery of warmed blood to the core during basking (Bartholomew, 1966; Seebacher and Franklin, 2003). In amphibians, however, the study of thermoregulation is complicated by high rates of evaporative cooling due to elevated cutaneous water loss. Indeed, amphibians have a relatively thin epidermis that allows for gas exchange (primarily oxygen uptake and carbon dioxide excretion) at the expense of high rates of cutaneous water loss (Feder and Burggren, 1985). Theoretical and empirical work suggest that amphibians prioritise the maintenance of water balance over thermoregulation (e.g., Anderson and Andrade, 2017; Tracy, 1976). Observations such as these led some to posit that responses to temperature in amphibians are mostly hydroregulatory instead of thermoregulatory (reviewed in Navas et al., 2008). However, amphibians have been shown to engage in behavioural thermoregulation both in the field (Lillywhite, 1970; Vences et al., 2002) and in the laboratory (Navas et al., 2021; Spotila, 1972). Moreover, some amphibians demonstrate active thermoregulation even in instances of obligate cutaneous respiration, as observed in lungless salamanders (Feder, 1982) and in frogs that overwinter under the ice of frozen lakes (Tattersall and Boutilier, 1997). Despite the evidence challenging the notion that amphibians may show little motivation to thermoregulate due to thermal constraints over desiccation (Carey, 1978), formal assessments of the behaviours involved in amphibian thermoregulation are lacking (Navas et al., 2021), especially in the context of seasonality.

Long-lived organisms experience seasonal shifts in climate multiple times over their lives (Williams et al., 2017), and seasonality has been shown to exert a strong selection pressure over the thermal biology of ectotherms (Clusella-Trullas et al., 2011; Giacometti et al., 2023). To cope with seasonality, some species may enter dormancy during periods of low environmental temperatures (Tattersall et al., 2016) or drought (Carvalho et al., 2010). While dormant, animals typically do not feed and remain mostly inactive to save energy until favourable climatic conditions return (Holden et al., 2021). Other species, however, may be naturally selected to express flexible phenotypes that aid with maintaining performance and thermal balance in the face of varying environmental conditions (Seebacher, 2005). In this case, seasonal changes in environmental parameters provide a signal for a shift in the thermal dependence of reaction norms (Seebacher et al., 2004) and thermoregulatory drive (sensu Navas et al., 2021), thereby eliciting acclimatisation. As a result, ectotherms from seasonal habitats typically exhibit different thermoregulatory behaviours, target different *T*_b_s between seasons, and are able to sustain activity across a wide range of *T*_b_s (Gilchrist, 1995; Hadamová and Gvoždík, 2011; Sepúlveda et al., 2008). Thus, by documenting seasonal changes in thermoregulatory parameters one may gain insight into the behavioural processes that underlie *T*_b_ control in ectotherms (Giacometti et al., 2023). Among vertebrate ectotherms, inquiry into the effect of seasonality over thermoregulation is taxonomically biased toward lizards (e.g., Clusella-Trullas and Chown, 2014; Giacometti et al., 2023; Huey and Pianka, 1977). Truly, despite the relatively large number of studies using amphibians as model systems in thermal biology research (e.g., Carilo Filho et al., 2022; Hutchison and Dupré, 1992), the extent to which thermal traits are capable of seasonal acclimation (i.e., response to a single environmental parameter) or acclimatisation (i.e., response to multiple environmental parameters) remains unclear (Navas et al., 2021; but see Pottier et al., 2022). Moreover, amphibian thermoregulation is known to vary extensively both inter- and intra-specifically, and also according to methodological procedures (e.g., acclimation temperatures, photoperiod regime) (Hutchison and Dupré, 1992; Navas et al., 2021).

Researchers investigating how seasonality affects thermoregulation in amphibians typically maintain animals under contrasting thermal and photoperiod regimes, and then allow the animals to make thermally-motivated choices within a laboratory thermal gradient (e.g., Feder, 1982; Heath, 1975). The rationale behind the use of thermal gradients to infer thermoregulation presupposes that thermoregulatory costs are low under laboratory conditions (Hertz et al., 1993), selected body temperatures (*T*_sel_) are an indication of the fundamental thermal niche of an organism (Magnuson et al., 1979), and movement within the gradient occurs due to thermotaxis (Doudoroff, 1938; sensu Navas et al., 2021). In amphibians, a change in *T*_sel_ is expected to be observed in response to seasonality. Studies in this group typically document seasonal changes in *T*_sel_ without specifically reporting on thermoregulatory behaviours (Navas et al., 2021). However, solely highlighting patterns of *T*_sel_ change does not provide enough information on the behavioural processes that underlie *T*_b_ control. Thus, it is important not to assume a priori that any seasonal change in *T*_sel_ occurs due to a shift in the motivation to thermoregulate (Giacometti et al., 2023), especially in species that may not use overt behaviours for *T*_b_ control. To overcome this limitation, behavioural observations may be integrated with *T*_sel_ measurements; past information on the study species is key to determine which behaviours should be scored (Giacometti et al., 2021). For instance, basking (Tattersall et al., 2006) or postural adjustments (Lillywhite, 1970) may be scored in assays with diurnal frogs, as these behaviours are associated with obtaining body heat via solar radiation in diurnal species from thermally variable habitats (Shine and Madsen, 1996). By contrast, microhabitat selection should be the main thermoregulatory behaviour in species from habitats where temperature is relatively stable (Catenazzi, 2016; Feder and Lynch, 1982), such as nocturnal and/or fossorial salamanders. For these species, assessing how movement patterns relate to changes in *T*_sel_ should be key to gauge seasonal shifts in the motivation to thermoregulate (Navas et al., 2021).

In this study, we evaluated the effect of seasonality over behavioural thermoregulation in the Spotted Salamander (*Ambystoma maculatum* Shaw, (1802)). Despite the large abundance of *A*. *maculatum* populations in parts of North America (Davis et al., 2018), the thermal biology of this species remains relatively understudied compared to other Caudata (e.g., Plethodontidae) (Feder, 1976; Spotila, 1972). The fossorial habit of adult *A*. *maculatum* suggests that this species spends most of its time within thermally stable environments, as underground refuges provide a thermal buffer to temperature extremes that may be experienced aboveground (Moore et al., 2018). Theory predicts that overt thermoregulatory behaviours should not be favoured in species from thermally stable habitats, given the limited thermoregulatory benefit (Ruibal, 1961; Shine and Madsen, 1996). Thus, one may expect that fossorial species should show relatively low evidence of thermotactic behaviours, and that seasonal fluctuations in temperature should not have a relevant impact over the thermal biology of these species (Giacometti et al., 2023; Wu et al., 2009). Importantly, although *A*. *maculatum* has a fossorial habit, its overall biology (e.g., reproduction, activity) follows well-delimited seasonal patterns (Sexton et al., 1990). Furthermore, a recent study demonstrated that body condition in a northern population of *A*. *maculatum* declined in response to increasingly warmer summer and autumn temperatures over the course of 12 years of monitoring (Moldowan et al., 2022). The same study also showed that *A*. *maculatum* experienced significant variation in underground temperatures across seasons (Moldowan et al., 2022). However, the extent to which seasonality influences behavioural thermoregulation in *A*. *maculatum* is still unclear.

We used an integrative approach to test how seasonal acclimatisation affected thermoregulatory behaviours within a thermal gradient. We first examined if salamanders showed different temporal patterns of *T*_sel_ between seasons. We tested for a position bias influencing thermoregulation by assessing whether salamanders that initially faced the cold end of the gradient would have lower *T*_sel_ compared to those that initially faced the warm end of the gradient (Wall and Shine, 2008). We then tested for a relationship between cutaneous water loss and *T*_sel_. We predicted that salamanders with high rates of cutaneous water loss would target relatively low *T*_sel_, as the selection of cool microhabitats minimises cutaneous water loss in amphibians (Brattstrom, 1979). We also tested whether salamanders traded increased locomotory activity for lower thermoregulatory precision. We predicted that highly mobile salamanders would sample a wider range of temperatures instead of targeting specific *T*_sel_ (Giacometti et al., 2021), thereby showing lower thermoregulatory precision than stationary salamanders. Besides quantifying conventional thermoregulatory parameters (Barber and Crawford, 1977), we also quantified *T*_sel_ skewness and kurtosis (DeWitt and Friedman, 1979) to further ascertain whether *A*. *maculatum* showed evidence of active thermoregulation. Lastly, we compared how thermoregulatory parameters differed between *A*. *maculatum* acclimatised to conditions that matched their active and overwintering seasons to test if salamanders showed greater evidence of thermophilic behaviours in the active season compared to the overwintering season (Feder and Lynch, 1982). Our study provides a broader understanding of thermoregulation in amphibians, particularly in the context of behavioural responses to seasonality in fossorial species.

## Material and methods

### Study site

Bat Lake is a naturally acidic (∼ 4.5–4.8 pH) and permanent kettle lake in Algonquin Provincial Park, Ontario, Canada (45.5857°N, 78.5185°W) (Bianchini et al., 2012). This lake is a fishless body of water, making it an ideal breeding ground for amphibians; predation by fishes is one of the main causes of amphibian egg mass mortality (Schmidt et al., 2021). Bat Lake is located at an altitude of ∼ 450 m above sea level, which is significantly higher than that of the surrounding low boreal wetland region (∼ 150 m above sea level) (Zoltai et al., 1988). From a climatic standpoint, this means that *A*. *maculatum* living in the coniferous forest surrounding Bat Lake experience climatic conditions comparable to those found in this species’ Northern range edge (Moldowan et al., 2022).

### Study species

*Ambystoma maculatum* (Caudata: Ambystomatidae) is a medium-sized species of mole salamander that has a wide distribution in eastern North America (O’Donnell, 1937). Despite being fossorial, this species is not a true burrower like others in its genus (Semlitsch, 1983). Instead, *A*. *maculatum* relies on burrows and tunnels built by small mammals (Regosin et al., 2003), but also dwells under rocks, coarse wood debris, and leaf litter (Faccio, 2003). Similar to other temperate zone ectotherms (Brodman and Jaskula, 2002), activity patterns in *A*. *maculatum* vary seasonally and are triggered by environmental cues, such as rainfall and temperature (Vasconcelos and Calhoun, 2004). In late winter/early spring, northern-latitude populations of *A*. *maculatum* emerge from underground burrows and migrate toward water bodies (e.g., ephemeral ponds or permanent lakes, in the case of our study population) to breed (Madison, 1997). This migratory period represents most of the aboveground activity in this species (Moldowan et al., 2022), although individuals may be observed foraging aboveground on relatively cool and rainy summer/autumn nights (Faccio, 2003). Based on long-term monitoring data, the approximate active period for the Bat Lake population of *A*. *maculatum* spans ordinal days 140–288 (May 20–October 17) (Moldowan et al., 2022).

On 15 May 2022, we collected 50 adult *A*. *maculatum* (26 females + 24 males) with the aid of a drift fence installed around the perimeter of Bat Lake. This drift fence was installed in the autumn of 2017 as part of the Bat Lake Inventory of Spotted Salamanders (BLISS) project (Moldowan, 2023). Logs and wood boards placed along the fence served as refuges for amphibians that reached the fence but were unable to cross it. Throughout the spring migratory period, researchers associated with BLISS conducted nighttime surveys to guarantee the timely processing and movement of amphibians. To ensure that all collected animals were at a similar physiological state (Finkler et al., 2003), we only collected post-breeding individuals, which we considered to be those that were found on the lake side of the drift fence (i.e., animals returning to the forest after breeding in the lake). We sexed individuals based on secondary sexual characteristics through visual inspection of cloacal morphology. During the breeding period, female *A*. *maculatum* have pleated cloacae, whereas males have papillose cloacae (Petranka, 1998).

We only collected animals that had not been marked and catalogued in previous BLISS monitoring efforts. When we collected an individual, we recorded its snout-vent length (SVL) with a tape measure, recorded its body mass with a Pesola™ spring scale (to the nearest 0.1 g), and assigned it a unique identification (ID); the ID was coupled with a photograph of the dorsum of the salamander to produce an individual-level recognition system based on spot patterns, as spots may vary in shape, size, and even colour in *A*. *maculatum* (Pierce and Shayevitz, 1982). Prior to transferring the salamanders to Brock University, we allocated them into plastic containers with ventilated fitted lids (34 cm x 19.6 cm x 12 cm). We kept a total of 10 individuals per container (totalling five containers during transport); each container had sphagnum moss, pine needles and water. We placed the containers inside a transport box kept at ∼ 4 °C so that animals would not risk overheating or dehydration during transportation. Prior to collecting salamanders, we obtained approval from Brock University’s Animal Care Committee (AUP #22-03-04), the Ministry of Northern Development, Mines, Natural Resources and Forestry (#1100575), and Ontario Parks.

### Husbandry

We housed salamanders in pairs within ventilated tanks that contained wet coconut husk fibre, sphagnum moss, and refuges made from PVC pipes; these tanks were stacked in racks to facilitate access to animals and misted regularly to keep local humidity high. We kept the housing tanks in a facility that had temperature, humidity, and photoperiod control. We changed temperature and photoperiod seasonally to mimic conditions experienced by *A*. *maculatum* in Bat Lake (Moldowan et al., 2022). Thus, our work represents an assessment of the effects of seasonal acclimatisation over thermal biology. In the spring, we kept salamanders at 7 °C, 70% relative humidity (RH), and a 10h:14h light:dark cycle. In the summer, we kept salamanders at 14 °C, 70% RH, and a 14h:12h light:dark cycle. In the autumn, we kept salamanders at 12 °C, 70% RH, and a 12h:12h light:dark cycle. In the winter, we kept salamanders at 2 °C, 70% RH, and under total darkness to simulate conditions inside overwintering burrows. To transition between spring to summer conditions, we increased temperature at the rate of 1 °C/day, totalling seven days necessary to go from 7 °C to 14 °C (from 14 July 2022 to 20 July 2022). The same protocol was used to transition between summer to autumn conditions, totalling two days needed to go from 14 °C to 12 °C (from 25 September 2022 to 26 September 2022). The transition between autumn to winter conditions followed a modified protocol, in which we decreased temperature at the rate of 1 °C every 2 ½ days, totalling 25 days necessary to go from 12 °C to 2 °C (from 17 November 2022 to 11 December 2022). Once a change in housing conditions was finalised, we always gave the salamanders at least four weeks to acclimatise to prevailing conditions before proceeding with our experiments. Throughout their period in the laboratory, we fed salamanders twice a week with mealworms dusted in calcium and multivitamin powder; water was available *ad libitum*. We also weighed salamanders to the nearest 0.01 g every week with an analytical scale (Mettler Toledo, model PB602-S) to monitor changes in body mass as a proxy for animal well-being.

### Thermal gradient set-up

To assess thermoregulation, we used an annular thermal gradient (total length = 120 cm, outer radius = 45 cm, inner radius = 30 cm) built by Brock University Technical Services (Fig. 1A). We purposely chose an annular thermal gradient to avoid low-temperature behaviours that could be mistaken with active temperature selection (e.g., “corner hugging”) (Giacometti et al., 2021); such behaviours are often observed in studies using rectangular thermal gradients, especially in fossorial species (e.g., Wall and Shine, 2008). The floor of the annular thermal gradient had copper pipes that were connected to hot- and cold-water baths (Haake™, models DC10 and SC100, respectively), which created a near linear temperature gradient (Fig. S1). For experiments conducted in the active season, the floor of the gradient ranged from ∼ 3 °C to 25 °C (range_active_ = 22 °C). In the overwintering season, the floor of the gradient ranged from ∼ −2.5 °C to 15 °C (range_overwintering_ = 17.5 °C). These temperature ranges represent possible habitat temperatures encountered by *A*. *maculatum* in Bat Lake during the active and overwintering seasons (Moldowan et al., 2022). We measured temperature along the thermal gradient to the nearest 0.01 °C with a thermocouple meter (Sable Systems, model TC-1000) before and after each experiment to ensure proper functioning of the gradient.

**Fig. 1.**
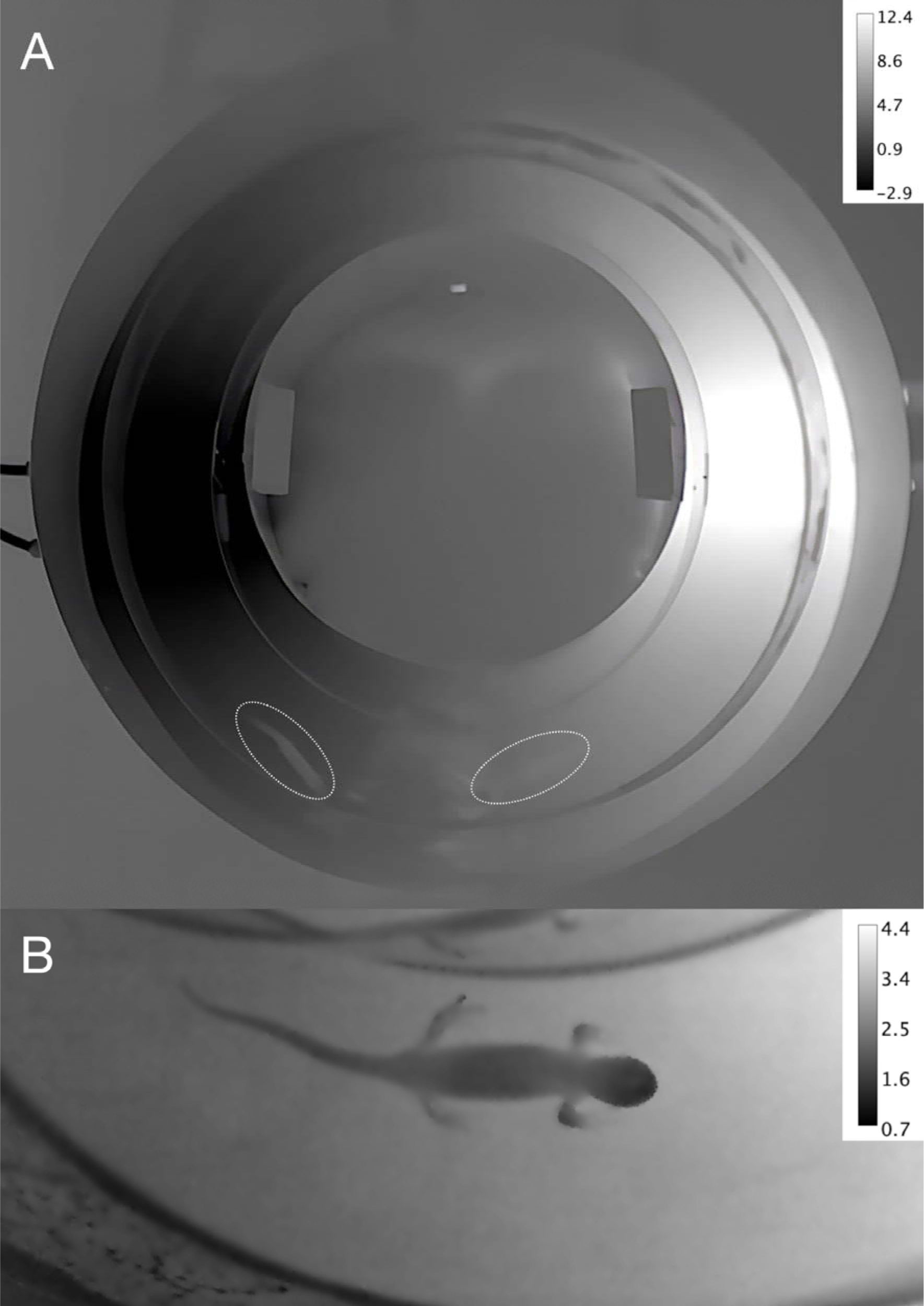
**A.** Thermal image of the annular thermal gradient used in this study depicting the temperature range used in the overwintering experiments. This thermal image was taken following the completion of an experiment, without the thermal gradient lid and partition being in view. The oval shapes denote two individuals of *Ambystoma maculatum*. The one on the lower left has a surface body temperature slightly higher than that of the floor of the thermal gradient. By contrast, the one on the lower right has a surface body temperature in equilibrium with that of the thermal gradient floor. **B**. Close-up thermal image of a female *A*. *maculatum* in the annular thermal gradient after the completion of an experiment. Despite the relatively small body size of the individual, one may still observe thermal heterogeneity among body parts. Note the different temperature range bars between panels **A** and **B**. In both panels, darker colours indicate lower temperatures and lighter colours indicate higher temperatures. Intermediate temperatures are shown in shades of grey.

The lid of the annular thermal gradient was affixed with a 10.5-cm tall partition that allowed the simultaneous study of two individuals per experimental run (i.e., one in the inner gradient lane, and another in the outer gradient lane). The lid of the thermal gradient also contained two venting holes every 90°, such that two venting holes were directly above the warm end at 0°, intermediate temperatures at 90° and 270°, and the cold end at 180°. Venting holes at 0°, 90°, and 270° were connected to a water vapour bubbling system put in place to minimise hydroregulation costs within the gradient. This system consisted of a 500-mL flask filled with dechlorinated water, and kept constantly at the warmest gradient temperature depending on the thermal condition (i.e., 25 °C in the active season or 15 °C in the overwintering season) to ensure the highest temperatures would experience high humidity. We put a 2.5-cm aquarium air stone inside the flask; the air stone was connected to a pressure pump (HIBLOW USA, model C-5BNS-0110) through a piece of tubing so that the water in the flask could be aerated. We sealed the flask with a rubber stopper that contained two openings in it: one for the tubing connecting the pressure pump to the air stone, and another to collect water vapour generated in the flask and direct it to the thermal gradient. The pressure pump was set to aerate the water in the flask at 1000 L/min, verified with a flow meter (Aalborg Instruments & Controls Inc., model PMR1-010972). The pressure created inside the flask directed water vapour from the flask to the venting holes in the gradient lid. We measured RH at different gradient temperatures with a data logger (MSR-Electronics GmbH, model MSR-145) to ensure that we had a continuous source of humid air at all gradient temperatures through the duration of our experiments. To avoid condensation due to water saturation in the gradient from limiting our ability to see the salamanders, we applied a water repellent (Stoner Invisible Glass®) to the interior of the thermal gradient lid before the start of each experiment. We used a paper towel to remove excess water repellent from the thermal gradient lid, so that the salamanders would not be in contact with excess liquid.

We set up the thermal gradient in the same facility where the salamanders were housed. We propped up the thermal gradient on 15 cm of insulation and foam padding to avoid building vibrations from disturbing the salamanders during experimentation. We conducted our experiments under complete darkness, matching the nocturnal habit of *A*. *maculatum*. We positioned two infrared illuminators (wavelength = 850 nm; TVPSii, model TP-IRBP15) 200 cm above the thermal gradient chamber to aid with visualising the animals through a livestream. To record the behaviour and position of salamanders inside the gradient, we fixed a high-resolution webcam (Agama, model V-1325R) 180 cm above the centre of the thermal gradient. This webcam was connected to a time lapse image acquisition software (HandyAVI®), which recorded an image every 30 s.

### Experimental design

The procedures described subsequently are applicable to the experiments conducted in both the active and overwintering seasons, unless otherwise stated. Importantly, in the current study, ‘active season’ pertains to salamanders acclimatised to summer conditions, and ‘overwintering season’ pertains to salamanders acclimatised to winter conditions. To measure behavioural thermoregulation, we allowed the salamanders (N = 50; randomly selected) a total of 8 h inside the thermal gradient (from ∼ 20h00 to 4h00 in the active season, and from ∼ 09h00 to 17h00 in the overwintering season). In both seasons, we gave the salamanders an initial 3-h habituation period, and used the data obtained in the subsequent 5 h (i.e., experimental period) in the analyses. We fasted salamanders for at least two weeks before experimentation to avoid thermophilic responses to feeding (Gvoždı k, 2003). We tested two individuals at a time, choosing at random whether a salamander would be tested in the inner or outer gradient lane. We always allowed a minimum interval of 12 h between trials after disinfecting the stage of the thermal gradient with 70% ethanol at the end of a trial.

Before introducing salamanders into the gradient, we placed them individually into a container filled with 30 mL of dechlorinated water for 15 min so water could be absorbed through the skin; we always handled the animals using nitrile gloves. Following this, we weighed the salamanders and placed them into the gradient, determining at random whether salamanders would be initially facing the cold or warm end of the gradient. For experiments done in the active season, we introduced the salamanders into the gradient at the 14 °C point, so that the first temperature they encountered therein would match their prevailing thermal condition. We followed the same protocol in the overwintering season, but at the 2 °C point. To minimise handling stress on thermoregulatory behaviours, we did not handle or manipulate the salamanders during the experiments.

Importantly, we verified that thermoregulatory differences (i.e., median *T*_sel_) between the active and overwintering seasons were not the outcome of offering different temperature ranges in the thermal gradient between seasons or conducting our experiments during a different time of day. To this end, we tested a subset of winter-acclimatised salamanders (N = 20) using a thermal gradient that ranged from ∼ −2.5 °C to 19.5 °C (hereafter “range control”; range_control_ = 22 °C, similar in span to the summer), and another subset of winter-acclimatised salamanders (N = 20) using the overwintering thermal gradient range from ∼20h00 to 4h00 (hereafter “time-of-day control”, conducted at the same time of day as the summer). After finishing the experiments, we removed the salamanders from the thermal gradient and measured their body mass. We used the difference in body mass before and after experiments as a proxy of cutaneous water loss during the 8 h of experimentation. After weighing, we placed the salamanders back into their housing tanks.

### Calculating T_sel_

To calculate *T*_sel_, we followed the protocol described in Giacometti et al. (2021). We captured a total of 960 images over the course of each 8-h long experiment, and imported the entire image sequence into the open-source image analysis software Fiji (Schindelin et al., 2012). From each image sequence, we recorded when the experiment was conducted (i.e., active or overwintering season), which individual was being tested in the inner and outer lanes, the sex of the individuals, and whether the salamanders were placed in the gradient facing the cold or warm end. We manually tracked the location of the salamanders in the thermal gradient for each image in the sequence using the manual tracking plug-in in Fiji (Schindelin et al., 2012). For each image sequence, we obtained a set of Cartesian (*x*, *y*) coordinates that had *x*, *y* = 0,0 as the lower left of the image. The following procedures describe how we used a series of calculations to convert salamander body position into selected gradient temperatures. Given that the centre of our images did not correspond to the centre of the thermal gradient (i.e., centroid), we subtracted the Cartesian coordinates of the centroid from those of the salamander’s position in the thermal gradient:

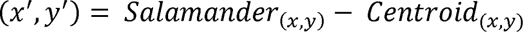

Where (*x*’, *y*’) represent normalised Cartesian coordinates. We converted these normalised Cartesian coordinates into distance by multiplying (*x*’, *y*’) values by 0.0010929 m/pixel; this conversion was determined empirically based on known gradient dimensions (see Giacometti et al., 2021). We then converted normalised Cartesian coordinates into polar coordinates (*r*, *θ*). Here, *r* is the hypothenuse determined by the *x*’ and *y*’ coordinates, and *θ* is the reciprocal of the *y*’/*x*’ tangent noted as degrees around a circle. For positive *y*’ values, *θ* ranged from 0° to 180°, and for negative *y*’ values, *θ* ranged from 0° to −180°. Given that the annular thermal gradient was symmetrical along the 0° horizontal axis, we converted *θ* values to an absolute value of |*θ|*. Lastly, we converted the final corrected *θ* position (*θ*’) into temperature. To do so, we used a second-order regression equation that described changes in temperature between known angular positions within the thermal gradient (Fig. S1). We empirically measured these changes in temperature along the gradient in all experiments.

In this experimental design, we assumed that the body temperature of the salamanders was in thermal equilibrium with the temperature of the gradient floor. Research in lizards (*Pogona vitticeps*, body mass range: 219–417 g) showed that *T*_sel_ matched core body temperatures over long time courses (< 0.5 °C difference) (Cadena and Tattersall, 2009). Considering the small body mass of the individuals used in our study (body mass range: 5.05–26.74 g) and the relatively long length of our experiments, we did not expect thermal inertia (Stevenson, 1985) to play a substantial role in heat exchange dynamics between the gradient floor and salamander body temperatures. That is, a relatively short time span would be necessary for salamander body temperature to equilibrate with gradient floor temperature. To verify this, we used a thermal camera (FLIR Systems, model FLIR T1030) to visually assess if the surface body temperature of an individual salamander matched the temperature of the thermal gradient. Assuming an object emissivity of 0.95 (Rowley and Alford, 2007), we processed the thermograph with ThermimageJ plugins for Fiji (Tattersall, 2019) that were verified against algorithms in FLIR Research Studio Software. We adjusted camera calibration constants and object parameters (e.g., relative humidity, object distance) according to conditions measured in the laboratory at the moment of thermal image sampling. To obtain measurements of dorsal mid-body (*T*_midbody_) and snout (*T*_snout_) skin temperature, we digitally drew a region of interest (ROI) over the salamander. We also drew an ROI over the thermal gradient floor immediately adjacent to the animal to obtain a measurement of gradient floor temperature (*T*_gradient_). From each ROI, we obtained an average temperature. We detected evidence of regional heterothermy (Webb et al., 1972) in the salamander, with the mid-point of the body showing average temperature values closest to those of the gradient floor compared to the snout (*T*_gradient_ = 3.16 °C; *T*_midbody_ = 2.46 °C; *T*_snout_ = 2.23 °C) (Fig. 1B). As such, we manually tracked the salamanders based on the position of the mid-point of their bodies, and used these values to calculate thermal set-point parameters (Barber and Crawford, 1977). Following image analysis, we partitioned each salamander’s surface body temperatures into percentiles. The 25^th^ percentile denotes the lower *T*_sel_ bound, the 50^th^ percentile denotes the median *T*_sel_, and the 75^th^ percentile denotes the upper *T*_sel_ bound. The difference between the upper and lower *T*_sel_ bounds is the *T*_sel_ range. These parameters form the bulk of the dual-limit system of body temperature control used to infer thermoregulatory drive in ectotherms. According to this framework, thermoregulatory drive should be minimal when body temperatures are within the lower and upper *T*_sel_ bounds (i.e., near median *T*_sel_). Positive thermotaxis (i.e., movement toward relatively warm temperatures) should be observed when animals are at body temperatures below the lower *T*_sel_ bound. In contrast, negative thermotaxis (i.e., movement away from relatively warm temperatures) should be observed when animals are at body temperatures above the upper *T*_sel_ bound. (Barber and Crawford, 1977; Barber and Crawford, 1979). Typically, animals that thermoregulate within narrow *T*_sel_ ranges are considered to be precise thermoregulators (DeWitt, 1967). In the context of seasonal shifts in thermal conditions, seasonal changes in *T*_sel_ parameters are interpreted as an indication of a species’ potential for plasticity in temperature-dependent traits (Giacometti et al., 2023).

### Total distance moved in the thermal gradient

To obtain a measure of activity within the gradient, we quantified the distance moved (*d*_t_) by each salamander every 30 s during the length of our experiments. Given that we used an annular thermal gradient, *d*_t_ follows the distance described by the arc of a circle:

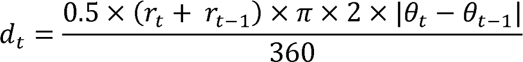

Where *r* is the mean radius, *θ* are polar coordinates, *t* and *t*-1 are adjacent time points for all *n* time points. To determine the total distance moved by each individual, we took the sum of *d*_t_:

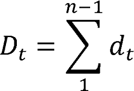

where *n* is the total number of images (*n* = 960).

### Data analysis

We performed all analyses using RStudio (version 2023.06.2) in R (version 4.2.2) (R Core Team, 2022) with a significance level of 0.05. We visually inspected Q-Q and P-P plots in search for deviations from normality and homoscedasticity using the ‘fitdist’ function from the fitdistrplus package (Delignette-Muller and Dutang, 2015). We built generalised linear models (GLMs) with the ‘glm’ function from the stats package (R Core Team, 2022), linear mixed-effects models (LMMs) with the ‘lmer’ function from the lme4 package (Bates et al., 2014), and generalised linear mixed-effects models (GLMMs) using the function ‘glmer’ from the lme4 package (Bates et al., 2014). To assess residual autocorrelation, we used the ‘checkresiduals’ function from the forecast package (Hyndman et al., 2020), and the ‘qqnorm’ and ‘acf’ functions from the stats package (R Core Team, 2022). We assessed model fit with the ‘check_model’ function from the performance package (Lüdecke et al., 2021). To visualise fixed model effects, we used the ‘plot’ function (graphics package; R Core Team, 2022) to plot an object created with the ‘allEffects’ function from the effects package (Fox et al., 2016). We created figures using the ggplot2 (Wickham, 2016), cowplot (Wilke et al., 2019), and patchwork (Pedersen, 2019) packages. All the data and code necessary to reproduce our analyses are available from Borealis through Brock University Dataverse at: https://doi.org/10.5683/SP3/X6BWUQ.

The following workflow was used to analyse data from both the active and overwintering seasons. To evaluate temporal changes in *T*_sel_ and determine the habituation and experimental periods, we fit a LMM that had median *T*_sel_ as the response variable, experimental time (binned into one-hour intervals) as the predictor, and salamander ID as the random term. Following this, we subset our *T*_sel_ data set and only analysed the experimental data. To improve the interpretation of model parameters, we used the ‘scale’ function (centre = TRUE, scale = FALSE) from the base package (R Core Team, 2022) to centre four variables: initial and final body mass, percentage of body mass loss, and total distance moved. We tested for differences in *T*_sel_ (response) between gradient lanes (predictor) using a GLM. Similarly, we used a GLMM to assess if total distance moved (response) differed between gradient lanes (predictor) while considering the pair of individuals tested in the inner and outer gradient lanes as a random term. Including a random term in this model resulted in a singular fit. Because of this, we built a GLM with total distance moved as the response variable and gradient lanes as the predictor, but without the random term. We calculated the Akaike Information Criterion (AIC) for each of these two models using the ‘AIC’ function from the stats package (R Core Team, 2022) and considered the model with the lowest AIC value to be the one with the best fit (Burnham and Anderson, 2002). To test for a difference between initial (pre-experiment) and final (post-experiment) body mass, we fit a LMM with body mass as the response variable, moment (i.e., initial or final) as the predictor, and salamander ID as a random term. We used a GLM to test for a relationship between body mass loss (%) and median *T*_sel_ while accounting for distance moved in the gradient and sex. We fit a GLM to test if the initial position that a salamander faced in the thermal gradient (i.e., cold versus warm) had an effect over median *T*_sel_ while accounting for body mass and sex. We built a GLM to assess if activity within the gradient affected thermoregulatory precision (i.e., *T*_sel_ range) while controlling for body mass and sex.

We assessed whether the skewness of individual *T*_sel_ distributions differed from zero using a one sample t-test fit with the ‘t.test’ function from the stats package (R Core Team, 2022). We then fit a GLM to test whether salamanders with higher *T*_sel_ (predictor) would show a more pronounced negative skew (response). To complement skewness observations, we also assessed the kurtosis of our individual-level *T*_sel_ data through a Jarque-Bera goodness of fit test using the ‘col_jarquebera’ function from the matrixTests package (Koncevičius, 2021). If kurtosis < 3, then the *T*_sel_ distribution is platykurtic, meaning that it produces fewer and less extreme outliers than a normal distribution. By contrast, if kurtosis > 3, then the *T*_sel_ distribution is leptokurtic, meaning that it produces more outliers than a normal distribution. From a biological perspective, a platykurtic *T*_sel_ distribution would indicate that salamanders are able to tolerate a wide range of gradient temperatures, whereas a leptokurtic *T*_sel_ distribution would point to salamanders targeting a narrow range of gradient temperatures.

Finally, we used GLMMs to compare whether thermoregulatory parameters, distance moved in the gradient, and body mass loss differed between the active and overwintering seasons. In all of these comparisons, we considered season as the predictor variable and salamander ID as a random term while also controlling for body mass and sex. As response variables, we considered median *T*_sel_, lower *T*_sel_ bound, upper *T*_sel_ bound, *T*_sel_ range, *T*_sel_ skewness, total distance moved, and body mass loss. We also fit GLMMs to test whether median *T*_sel_, lower *T*_sel_ bound, upper *T*_sel_ bound, *T*_sel_ range, and total distance moved differed between the overwintering season and the range control, and between the overwintering season and the time-of-day control. We performed Bonferroni correction of p-values when multiple models were fit on data derived from the same subset of experiments through the ‘p.adjust’ function from the stats package (R Core Team, 2022). Detail on model families, fixed effects, and random effects can be found in Table S1.

## Results

### Temporal trends in T_sel_ and validation of experimental assumptions

In both seasons, we found that *T*_sel_ varied over the 8 h of experiment (active: χ^2^ = 19.60, *p* = 0.01; overwintering: χ^2^_(8)_ = 334.03, *p* = 2.2×10^−16^). Median *T*_sel_ values were consistently offset higher than prevailing thermal conditions in both seasons (Fig. 2). Median *T*_sel_ did not differ between the inner and outer gradient lanes in neither the active (χ^2^_(1)_ = 0.46, *p* = 0.49) or overwintering seasons (χ^2^_(1)_ = 0.67, *p* = 0.41). In both seasons, we obtained singular fits in models evaluating if total distance moved differed between gradient lanes while considering ‘Pair’ as a random term. Comparisons of AIC values between LMMs and GLMs indicated GLMs to be of better fit (see Code in the Supplementary Material). Based on GLMs, we found that total distance moved was similar between lanes in the active (χ^2^_(1)_ = 0.10, *p* = 0.76) and overwintering seasons (χ^2^_(1)_ = 0.01, *p* = 0.94). Body mass differed between the beginning and the end of the experiments (active: χ^2^ = 168.51, *p* < 2.2×10^−16^; overwintering: χ^2^ = 77.26, *p* < 2.2×10^−16^). Salamanders lost, on average, 0.15 g/h in the active season compared to 0.04 g/h in the overwintering season during the 8-h long experiments. Despite this, body mass loss was not influenced by median *T*_sel_, distance moved, or sex within either season (see code in the Supplementary Material). Our water vapour bubbling system kept high RH values at all gradient temperatures ranging from −2.5 °C to 25 °C (average ± standard deviation = 92.93 ± 6.47% RH).

**Fig. 2.**
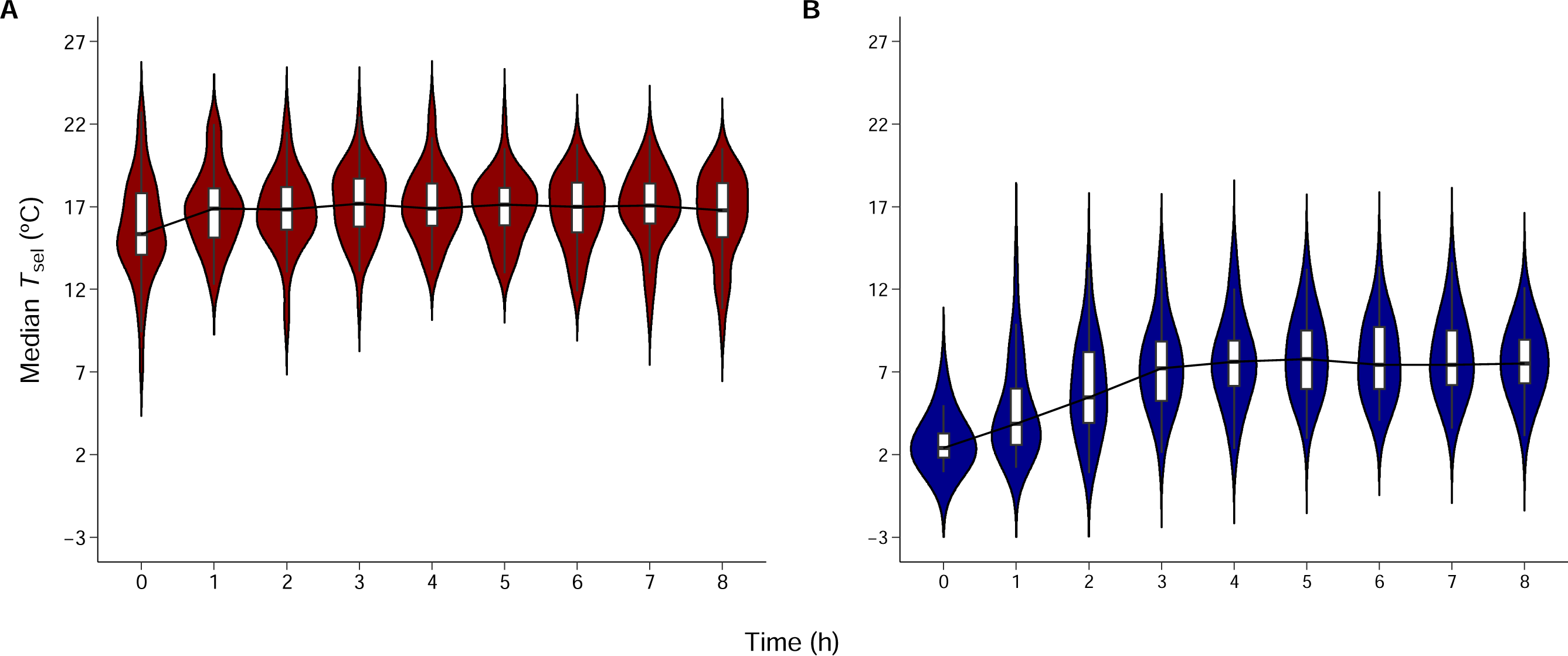
Temporal variation in the selected body temperature (*T*_sel_) of *Ambystoma maculatum* throughout 8 h in an annular thermal gradient during the (**A**) active and (**B**) overwintering seasons. In both seasons, we binned the 8 h of data into 1-h intervals. In both panels, the grey dashed lines represent the acclimatisation temperatures in the active (14 °C) and overwintering (2 °C) seasons. Boxes denote the among-individual lower (25^th^) and upper (75^th^) bounds of *T*_sel_ at a given hour. The horizonal line in each box indicates the median *T*_sel_, and the black line connecting each median *T*_sel_ showcases the hourly change in this parameter. Temperatures selected by each individual salamander are shown as semi-transparent dots. The violin plot shows the kernel density probability of *T*_sel_ data at different hours of experimentation.

### Behavioural thermoregulation parameters

We used the experimental period to calculate the median, lower, and upper *T*_sel_ bounds from individual animal *T*_sel_ distributions (Table 1). In the active season, we determined *A*. *maculatum* to have a median *T*_sel_ of 17.00 °C, a lower *T*_sel_ bound of 12.60 °C, and an upper *T*_sel_ bound of 20.10 °C (Fig. 3). In the overwintering season, *A*. *maculatum* had a median *T*_sel_ of 7.61 °C, a lower *T*_sel_ bound of 4.89 °C, and an upper *T*_sel_ bound of 10.40 °C (Fig. 3). Whether salamanders were initially facing the cold or warm end of the gradient had no effect over median *T*_sel_ in either season; sex and body mass also did not affect median *T*_sel_ (Table S2).

**Fig. 3.**
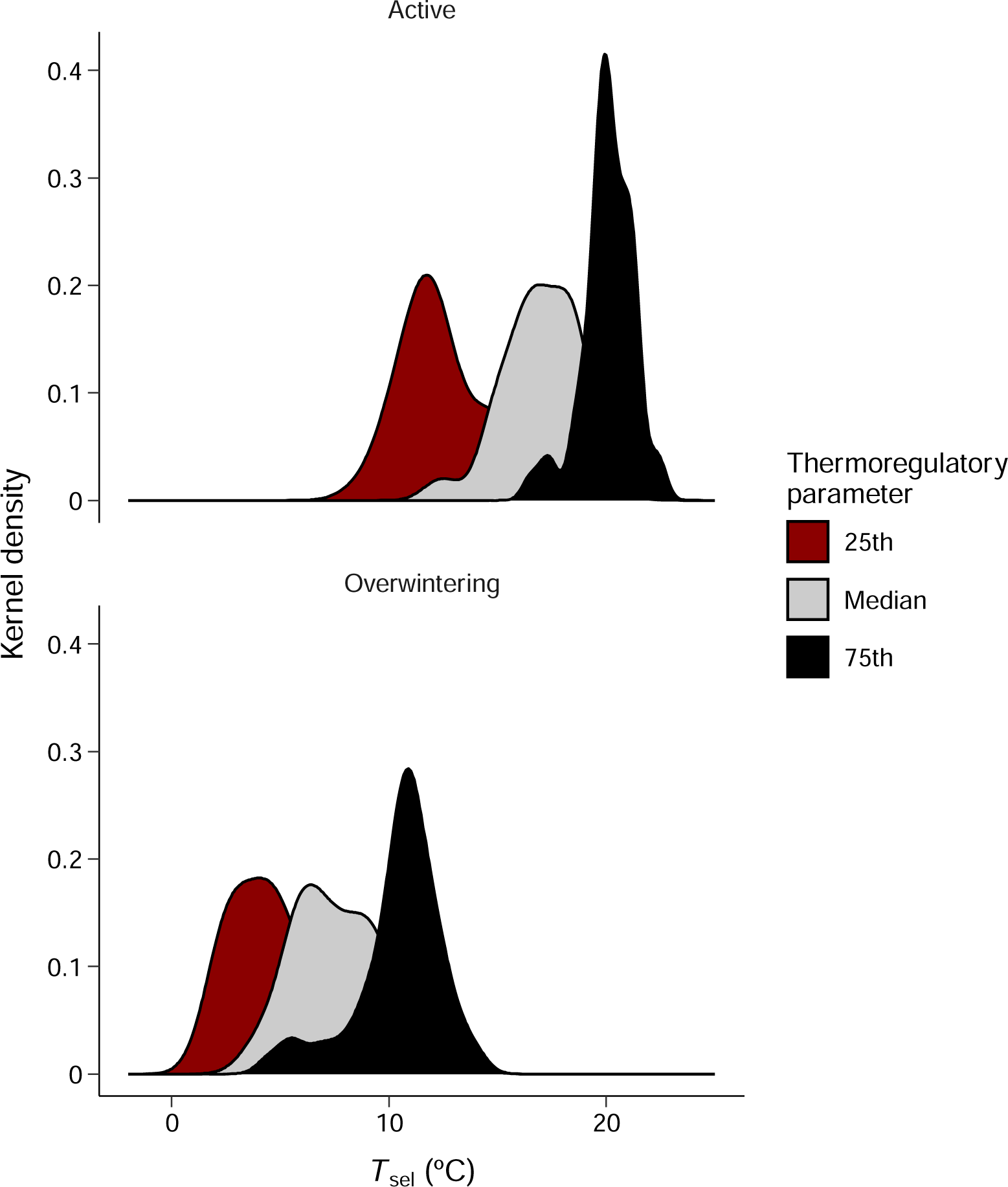
Kernel density estimations of individual level temperature selection (*T*_sel_) parameters for *Ambystoma maculatum* in the active and overwintering seasons. Parameters are colour-coded, such that red represents the lower (25^th^) *T*_sel_ bound, grey denotes the median *T*_sel_, and black indicates the upper (75^th^) *T*_sel_ bound.

**Table 1.** Parameters associated with temperature selection (*T*_sel_) and behavioural thermoregulation for *Ambystoma maculatum* studied in the active and overwintering seasons. *p*-values in bold denote significant differences in a given trait between the active and overwintering seasons. SD = standard deviation; Min = minimum value; Max = maximum value; χ^2^= chi-square.

|  | Active<br>(N=50) | Overwintering<br>(N=50) | Comparisons |
| --- | --- | --- | --- |
| **Median $T_{\text{sel}}$ (°C)** |  |  |  |
| Mean ± SD | 17.00 ± 1.77 | 7.61 ± 2.10 | $\chi^2_{(1)} = 734.00, p < \mathbf{0.001}$ |
| Median (Min, Max) | 17.40 (12.10, 20.60) | 7.37 (3.83, 13.20) |  |
| **Lower $T_{\text{sel}}$ bound (°C)** |  |  |  |
| Mean ± SD | 12.60 ± 2.11 | 4.89 ± 2.60 | $\chi^2_{(1)} = 263.42, p < \mathbf{0.001}$ |
| Median (Min, Max) | 12.40 (8.22, 20.10) | 4.46 (1.78, 13.20) |  |
| **Upper $T_{\text{sel}}$ bound (°C)** |  |  |  |
| Mean ± SD | 20.10 ± 1.15 | 10.40 ± 2.00 | $\chi^2_{(1)} = 1022.76, p < \mathbf{0.001}$ |
| Median (Min, Max) | 20.30 (17.20, 22.40) | 10.80 (4.49, 14.10) |  |
| **$T_{\text{sel}}$ range (°C)** |  |  |  |
| Mean ± SD | 7.48 ± 1.91 | 5.53 ± 2.65 | $\chi^2_{(1)} = 18.19, p < \mathbf{0.001}$ |
| Median (Min, Max) | 7.85 (0.80, 11.20) | 6.18 (0.07, 9.16) |  |
| **Skewness** |  |  |  |
| Mean ± SD | -0.51 ± 0.44 | -0.38 ± 0.97 | $\chi^2_{(1)} = 0.81, p = 0.36$ |
| Median (Min, Max) | -0.53 (-2.09, 0.80) | -0.19 (-5.41, 1.10) |  |
| **Total distance moved (m)** |  |  |  |
| Mean ± SD | 109.00 ± 67.10 | 78.20 ± 70.50 | $\chi^2_{(1)} = 7.10, p = 0.054$ |
| Median (Min, Max) | 95.20 (0.26, 336) | 58.00 (0.14, 262) |  |
| **Body mass loss (%)** |  |  |  |
| Mean ± SD | 8.68 ± 3.40 | 2.66 ± 1.84 | $\chi^2_{(1)} = 49722.03, p < \mathbf{0.001}$ |
| Median (Min, Max) | 8.95 (0.20, 14.7) | 2.18 (0.11, 7.37) |  |

Note: we made comparisons between seasons using linear mixed effects models that had season, body mass, and sex as the predictor variables, as well as salamander ID as a random term. Given that neither body mass or sex mediated differences between seasons, we only report $\chi^2$ and $p$ -values for season in each model.

To assess how activity in the gradient affected thermoregulatory precision, we examined the relationship between *T*_sel_ range and total distance moved while accounting for sex and body mass effects. In both seasons, we found that the more an individual moved, the wider its *T*_sel_ range (i.e., the lower its thermoregulatory precision). Neither sex nor body mass mediated this effect (Table 2). In the active season, the individual distribution of *T*_sel_ was negatively skewed and different from a normal distribution (skewness_active_ = −0.50; *t* = −8.24, *p* = 8.28×10^−11^). The Jarque-Bera goodness of fit test indicated that *T*_sel_ was platykurtic in the active season (kurtosis = 2.15) (Fig. S2). We obtained similar results for skewness (skewness_overwintering_ = −0.38; *t* = −2.74, *p* = 0.01) and kurtosis (kurtosis = 2.07) in the overwintering season (Fig. S2). In both seasons, individuals with higher median *T*_sel_ showed more negative skew (β_active_ = −−0.20; *t* = −7.41, *p* = 2×10^−9^; β_overwintering_ = −0.30; *t* = −6.22, *p* = 1.14×10^−7^; Fig. 4).

**Table 2.** Parameter estimates (β), 95% confidence intervals (95% CI), and *p*-values for the models assessing how total distanced moved affected thermoregulatory precision in *Ambystoma maculatum*. The models had *T*_sel_ range as the response variable, distance moved, body mass, and sex as the predictor variables. Significant parameters are denoted in bold.

| Active |  |  |  |
| --- | --- | --- | --- |
| *Predictors* | $\beta$ | 95% CI | $p$ |
| Intercept | 7.91 | 7.22 – 8.61 | **<0.001** |
| Total distance moved | 0.02 | 0.01 – 0.02 | **<0.001** |
| Sex (Male) | 0.05 | -0.10 – 0.21 | 0.492 |
| Body mass | -0.89 | -2.02 – 0.23 | 0.120 |
| Observations | 50 |  |  |
| R<sup>2</sup> | 0.368 |  |  |
| Overwintering |  |  |  |
| *Predictors* | $\beta$ | 95% CI | $p$ |
| Intercept | 5.18 | 4.40 – 5.97 | **<0.001** |
| Total distance moved | 0.03 | 0.02 – 0.04 | **<0.001** |
| Sex (Male) | 0.03 | -0.12 – 0.18 | 0.679 |
| Body mass | 0.73 | -0.56 – 2.02 | 0.270 |
| Observations | 50 |  |  |
| R<sup>2</sup> | 0.590 |  |  |

**Fig. 4.**
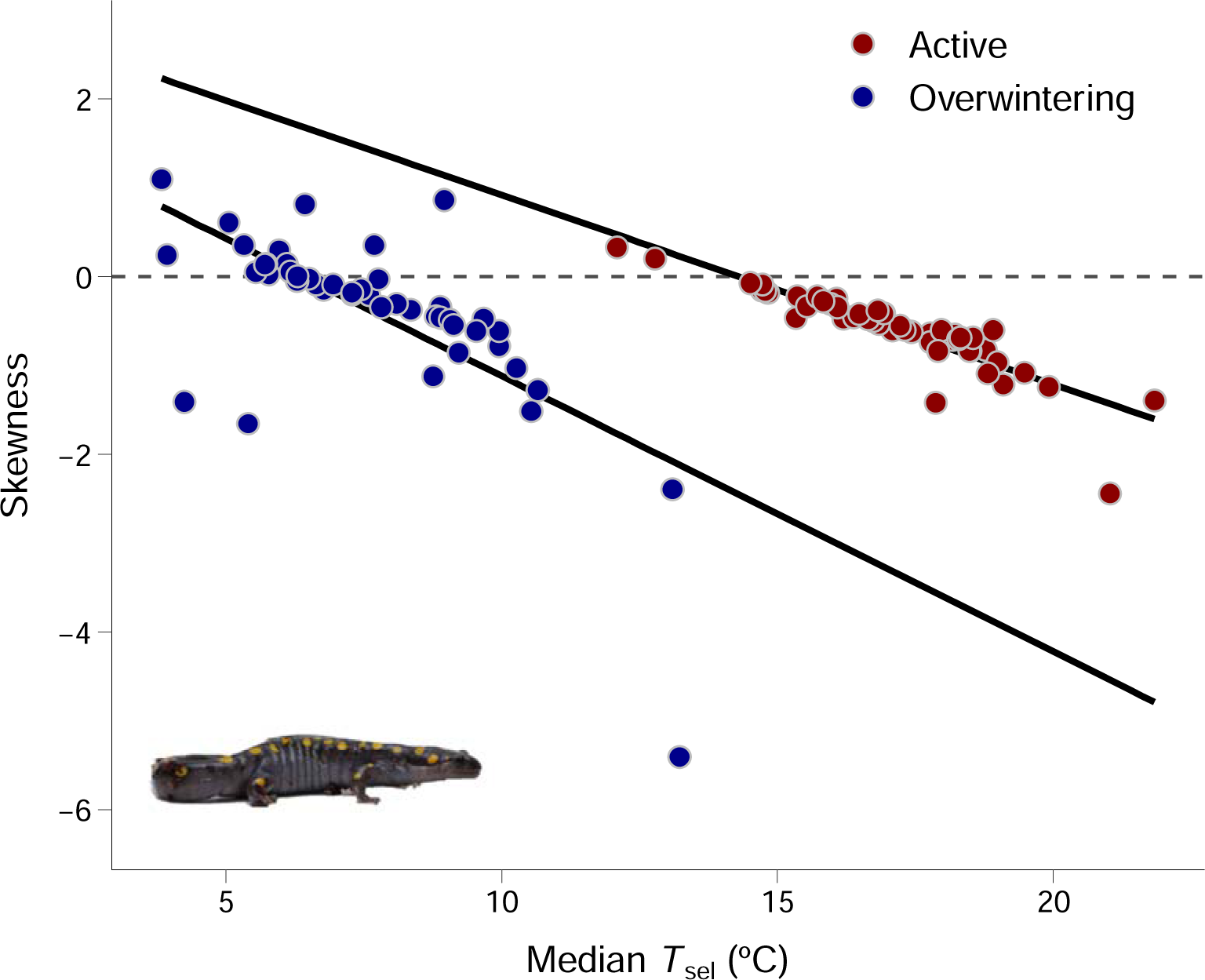
Relationship between skewness and median selected temperatures (*T*_sel_) for *Ambystoma maculatum* in the active (red) and overwintering (blue) seasons. The solid black lines and shaded areas indicate the predicted relationship between skewness and *T*_sel_, and the 95% confidence interval, respectively. The grey dashed line indicates the zero. Colour-coded dots represent individual *A*. *maculatum* (N = 50 per season). The inset shows a female *A*. *maculatum*.

All thermoregulatory parameters studied herein differed between seasons, except for skewness and total distance moved, with no effect of sex or body mass (Table 1). We did not find thermoregulatory differences between animals tested in the overwintering season and the range control, except for *T*_sel_ range and total distance moved (Table S3). Similarly, we did not find thermoregulatory differences between salamanders tested in the overwintering season and the time-of-day control, except for *T*_sel_ range and total distance moved (Table S4).

## Discussion

Here, we evaluated the effect of seasonal acclimatisation over *T*_sel_ and behavioural thermoregulation in *A*. *maculatum*. Adult *A*. *maculatum* live in underground burrows that provide a thermal buffer against shifts in aboveground temperatures (Moldowan et al., 2022; Moore et al., 2018). Theory predicts relaxed selection pressure over thermoregulation in species from thermally stable habitats (e.g., fossorial or tropical species) (Greer, 1980; Shine and Madsen, 1996). As a result, these species typically have relatively low *T*_sel_, low thermoregulatory precision, and show low evidence of thermophilic behaviours compared to species from thermally variable habitats (Camacho et al., 2015; Withers, 1981). While these patterns are well established in reptiles, we still have a limited understanding of amphibian thermoregulation in the context of fossoriality. By documenting how thermal biology traits shift in response to seasonal acclimatisation, we provide evidence that adult *A*. *maculatum* engage in active behavioural thermoregulation despite being fossorial.

### The effect of experimental design over inferences of behavioural thermoregulation

We found that *T*_sel_ varied over the course of our 8-h long experiments, which was consistent with studies evaluating hourly changes in the *T*_sel_ of amphibians (Heath, 1975; Hutchison and Spriestersbach, 1986). Although daily shifts in *T*_sel_ may be regulated by endogenous aspects (e.g., thyroid hormones) (Zena et al., 2020), the pattern we observed is more likely explained by animals gaining a spatial awareness of the thermal gradient (Cadena and Tattersall, 2009) and recovering from stress caused by manipulation (Cabanac and Gosselin, 1993) in the initial hours of the experiment. Our study also validated assumptions relevant to the design of behavioural thermoregulation experiments. Measuring the *T*_sel_ of multiple individuals at a time is common practice in thermal ecology research (Gaudenti et al., 2021; Gilbert and Miles, 2019). Researchers typically introduce animals into separate gradient lanes and assume that thermoregulation is not affected by experimental design. This, however, is rarely verified (Spotila, 1972), and overlooks the possibility of animals altering their thermoregulatory behaviour from detecting the presence of nearby organisms. For example, *Amalosia lesueurii* (Duméril & Bibron, 1836) (Squamata: Diplodactylidae) were shown to avoid thermally suitable areas after recognising the scent of potential predators (Webb et al., 2010). Moreover, *Salamandra salamandra* (Linnaeus, 1758) (Caudata: Salamandridae) were shown to be capable of sex discrimination through olfaction, which resulted in individuals changing their spatial distribution after being presented with the scent of a conspecific of the opposite sex (Caspers and Steinfartz, 2011). Whether the ability of amphibians to sense a nearby conspecific has an impact over realised thermoregulation is unknown.

In both seasons, we found that median *T*_sel_ and total distance moved were similar between *A*. *maculatum* tested in the inner and outer gradient lanes. Thus, our data suggest that salamanders made use of thermoregulatory behaviours regardless of the presence of a nearby conspecific. Different patterns of thermoregulation between lanes could be indicative of *A*. *maculatum* avoiding specific parts of the gradient after detecting a conspecific through olfaction, which was the main sensory cue available in our experimental design. Although olfaction is a key orientation mechanism in nocturnal species (McGregor and Teska, 1989), amphibians integrate different stimuli into a multisensory system to obtain directional information (Sinsch, 2006). To date, studies have shown that the ability of tree frogs to change skin colour in response to visually perceived changes in background brightness has thermoregulatory consequences (King et al., 1994; Tattersall et al., 2006). However, our understanding of how other sensory stimuli (e.g., acoustic, magnetic, mechanic) may interact with thermal biology is still limited and warrants further research.

Work in the in the fossorial *Lialis burtonis* Gray, 1835 (Squamata: Pygopodidae) suggested that individuals would end up with relatively low *T*_sel_ if introduced into the cold end of a thermal gradient; the opposite was true when lizards were introduced into the warm end (Wall and Shine, 2008). These data suggested that *L*. *burtonis* tolerated rather than actively selected gradient temperatures, which corroborated the expectation of low evidence of thermotactic behaviours in fossorial ectotherms. In our study, we controlled for the location at which *A*. *maculatum* were first placed in the thermal gradient by always introducing them at the temperature that corresponded to the prevailing acclimatisation regime. The initial direction *A*. *maculatum* faced when introduced into the thermal gradient did not impact median *T*_sel_ in either season. Thermal gradient design could explain why a position bias over *T*_sel_ was observed in *L*. *burtonis* but not in *A*. *maculatum* despite both species being fossorial. Indeed, *L*. *burtonis* were tested in a rectangular gradient (Wall and Shine, 2008), whereas we tested *A*. *maculatum* in an annular thermal gradient. Rectangular thermal gradients are the most common gradient type in thermal biology studies (e.g., Camacho and Rusch, 2017; Hertz et al., 1993; Navas et al., 2021). However, the rectangular shape may produce a site-specific bias towards ends or corners of the thermal gradient, resulting in misleading inferences of *T*_sel_ (Reiser et al., 2013). Although less used in thermal biology research (but see Bowker, 1984; DeWitt, 1967), annular thermal gradients are argued to allow for easier navigation (Giacometti et al., 2021). The absence of corners in annular thermal gradients also encourages animals to actively select gradient temperatures (Reiser et al., 2013), thereby facilitating assessments of thermotaxis and the relationship between locomotion and *T*_sel_. Importantly, formal assessments of whether rectangular and annular gradients differ in the ability to infer thermoregulation have yet to be made in terrestrial ectotherms.

We found that body mass differed between the start and the end of the experiments in both seasons. However, body mass loss was not explained by median *T*_sel_, total distance moved, or sex. Thus, salamanders with relatively high body mass loss did not move toward cooler areas of the gradient as a primary defense mechanism against possible desiccation (Brattstrom, 1979). This suggests that we were able to minimise hydroregulatory costs within the thermal gradient, and that our experimental conditions did not compromise salamander hydration status, which is a critical factor in amphibian thermoregulation (Ortega et al., 2023; Tracy et al., 1993). Indeed, the lowest RH value created through our water vapour bubbling system was ∼ 80% at 12 °C in the active season. While information on soil moisture preference is unavailable for *A*. *maculatum*, non-breeding individuals of the congener *A*. *opacum* (Gravenhorst, 1807) preferred soils of 74% RH at 13 °C (Marangio and Anderson, 1977). Coupled with the fact that we never observed our study individuals employing hydroregulatory behaviours (e.g., coiling) (Heatwole, 1960), these results suggest that our behavioural observations were thermoregulatory in nature. To obtain a broader understanding of the trade-off between thermoregulation and hydroregulation in amphibians, future work may assess whether *A*. *maculatum* uses behavioural adjustments to defend a preferred RH range once thermoregulatory costs are minimised.

### Seasonal acclimatisation of behavioural thermoregulation

We found that *A*. *maculatum* traded off increased locomotion for lower thermoregulatory precision, as evidenced by a positive relationship between *T*_sel_ range and total distance moved. This effect was consistent in salamanders tested in both seasons, and matches a pattern previously reported in the semi-fossorial *Storeria occipitomaculata occipitomaculata* (Storer, 1839) (Squamata: Colubridae) (Giacometti et al., 2021). Additionally, this trade-off was not mediated by sex or body mass in neither season, suggesting a possible link between activity levels and thermoregulation irrespective of sexual size dimorphism (Bulté and Blouin-Demers, 2010). Importantly, movement within a laboratory thermal gradient is primarily associated with active temperature selection. In nature, however, movement patterns may depend on factors such as foraging tactics, the presence of nearby predators, and climate (Hileman and Brodie Jr, 1994; Jaeger and Barnard, 1981; O’Donnell et al., 2016), all of which may conflict with an ectotherm’s motivation to thermoregulate (Huey and Slatkin, 1976). A recent mark-recapture study pointed to a link between minimum temperatures and increased movement in non-breeding *A*. *maculatum*, *A*. *jeffersonianum* (Green, 1827), and *A*. *laterale*-*jeffersonianum* Hallowell, 1856 (Van Drunen et al., 2024). However, the interplay between salamander movement and thermoregulation in natural settings is still unclear. By combining laboratory inferences of thermoregulation along with field measurements of *T*_b_ and radio-tracking techniques (e.g., Forget-Klein and Green, 2021; Tattersall et al., 2016), one may clarify whether behavioural thermoregulation is relevant to salamanders in nature by evaluating how movement contributes to the maintenance of thermal balance.

Our data on *T*_sel_ skewness and kurtosis supported the prediction that ectotherms should be more tolerant of low temperatures than high temperatures (Martin and Huey, 2008). Indeed, in both seasons we found individual *T*_sel_ to be negatively skewed and platykurtic, which fits the general characteristics of thermal performance curves, especially in thermal generalists (Gilchrist, 1995; Huey and Pianka, 2018). We also found that *A*. *maculatum* with higher *T*_sel_ showed a more pronounced negative skew than those with lower *T*_sel_ in both seasons. The reasons behind the ubiquity of the negative skew in ectotherm *T*_b_ and *T*_sel_ are likely manyfold (see Huey and Pianka, 2018). From a thermosensory standpoint, warm-sensitive neurons are more responsive than cold-sensitive ones (Cabanac et al., 1967), which may result in enhanced sensory-driven thermal behaviours at relatively warm temperatures. Alternatively, the negative skew in *T*_sel_ could be a product of the exponential relationship between *T*_b_ and physiological performance (DeWitt and Friedman, 1979). Under this view, performance would be reduced at relatively low temperatures and increased at relatively high temperatures in species that possess asymmetric and narrow thermal performance curves (Huey and Pianka, 2018). However, ectotherms from highly seasonal habitats tend to fit the description of thermal generalists, which are animals that trade off performance peak for a wide performance breadth (Gilchrist, 1995). Such species often show seasonal lability in thermal biology traits, which enhances the maintenance of thermal balance across seasons (Giacometti et al., 2023). Quantifying seasonal changes in performance traits should clarify if *A*. *maculatum* is a thermal generalist and further our knowledge of the possible drivers of the negative *T*_sel_ skew in ectotherms.

Our results supported the presence of active thermoregulation in *A*. *maculatum*, highlighting that seasonal acclimatisation prompted a shift in *T*_sel_ and associated thermotactic behaviours. However, thermoregulation in *A*. *maculatum* may not be as precise as in species that live primarily aboveground. Indeed, we found that *A*. *maculatum* consistently thermoregulated between relatively wide *T*_sel_ ranges. The *T*_sel_ range value we obtained in the active season is comparable to that of the fossorial *Saiphos equalis* (Gray, 1825) (Squamata: Scincidae), which had a mean *T*_sel_ range of 7 °C (Wu et al., 2009). Our *T*_sel_ values are also consistent with the range of values reported in laboratory studies of amphibian *T*_sel_ (reviewed in Hutchison and Dupré, 1992). For instance, our *T*_sel_ measurements in the active season are similar to those of summer-caught *A*. *tigrinum* (Green, 1825) (mean ± standard deviation; *T*_sel_ = 18.4 ± 0.6 °C), a fossorial congener of comparable body size (Heath, 1975). We also found seasonal differences in all thermoregulatory parameters except for *T*_sel_ skewness. These results are supported by previous research that indicated a heightened acclimatory capacity of *T*_sel_ and critical thermal maximum in *A*. *maculatum* relative to *A*. *texanum* Matthes, 1855 (Keen and Schroeder, 1975). Our analyses indicated that *A*. *maculatum* thermoregulated more precisely in the overwintering season than in the active season. This is a recurrent pattern in lizards, typically interpreted as increased energetic allocation toward thermoregulation in the face of adverse climatic conditions (Hertz et al., 1993; Patterson and Davies, 1978). In our study, however, increased thermoregulatory precision in the overwintering season may be explained by an effect of seasonal acclimatisation over locomotion. Winter-acclimatised *A*. *maculatum* decreased their locomotory activity compared to the experiments conducted in the summer, which may have translated into *A*. *maculatum* thermoregulating between a narrower *T*_sel_ range in the overwintering season compared to the active season. This observation may also explain why the *T*_sel_ range and total distance moved differed between the range control and overwintering experiments, as well as why the *T*_sel_ range and total distance moved differed between the time-of-day control and overwintering experiments.

## Conclusions

The cryptic nature of many fossorial ectotherms has limited research into their physiology and behaviour (Jared et al., 1999). This resulted not only in knowledge gaps about their basic biology, but also in an overreliance over untested assumptions about the role of fossoriality in shaping the physiology and behaviour of ectotherms (Giacometti and Tattersall, 2023). Our study highlights how seasonal acclimatisation affects behavioural thermoregulation in the fossorial *A*. *maculatum* through seasonal changes in the use of thermal resources. Our work challenges the expectation of low evidence of thermophilic behaviours in fossorial species by showing *T*_sel_ values to be consistently offset higher than prevailing thermal conditions in both the active and overwintering seasons. Moreover, our finding of a 9.4-°C shift in *T*_sel_ between seasons adds to mounting evidence in support of seasonal lability in thermal traits (e.g., Clusella-Trullas and Chown, 2014; Giacometti et al., 2023), and against the use of the “final thermal preferendum” concept as a unifying paradigm in ectotherm thermal biology (Reynolds and Casterlin, 1979). We emphasise the importance of integrating behavioural observations with measurements of thermal biology traits to better understand the mechanisms underlying *T*_b_ control in amphibians (Navas et al., 2021). We also highlight that considering the influence of seasonality over thermal biology should contribute to the goal of elucidating behavioural responses to environmental variability in ectotherms (Giacometti et al., 2023). Ultimately, our findings are aligned with recent work showing that temperature seasonality is a relevant aspect explaining a long-term shift in body size and condition in *A*. *maculatum* (Moldowan et al., 2022). Given that thermal biology underpins most organismal processes in ectotherms, an inability to realise *T*_sel_ in the field could translate into salamanders being unable to maximise energy turnover and thus explain why *A*. *maculatum* are getting smaller over time. In this context, future work may undertake an energetic perspective, and assess how energy metabolism varies in response to seasonal shifts in temperature.

## Acknowledgements

We thank Ontario Parks, the Ministry of Northern Development, Mines, Natural Resources and Forestry, the Algonquin Wildlife Research Station, Njal Rollinson, Kevin Kemmish, and Patrick Moldowan for facilitating access to Bat Lake and our study animals. We are grateful to all the researchers who were ever involved with the Bat Lake Inventory of Spotted Salamanders (BLISS). We thank Shawn Bukovac, Kristin Bray, Tom Eles, Sarah Kehoe, Kate Luttjehuizen, Jordan Albanese, Ahmad Mohammad, and Natasha Hearn for their assistance with animal care. We extend our thanks to Stephen Renda, Mitch Sillaste, and Art Reimer from Brock University’s Machine Shop for building the thermal gradient used in our study.

## Author contributions

DG and GJT conceived the idea of the study. GJT acquired research funding. DG performed the experiments. DG analysed the data with input from GJT. DG drafted the manuscript. DG and GJT contributed to manuscript edits and revisions.

## Data accessibility statement

The data and code necessary to reproduce our analyses can be accessed from: https://doi.org/10.5683/SP3/X6BWUQ.

## Supplementary material

### Supplementary tables

**Table S1.**
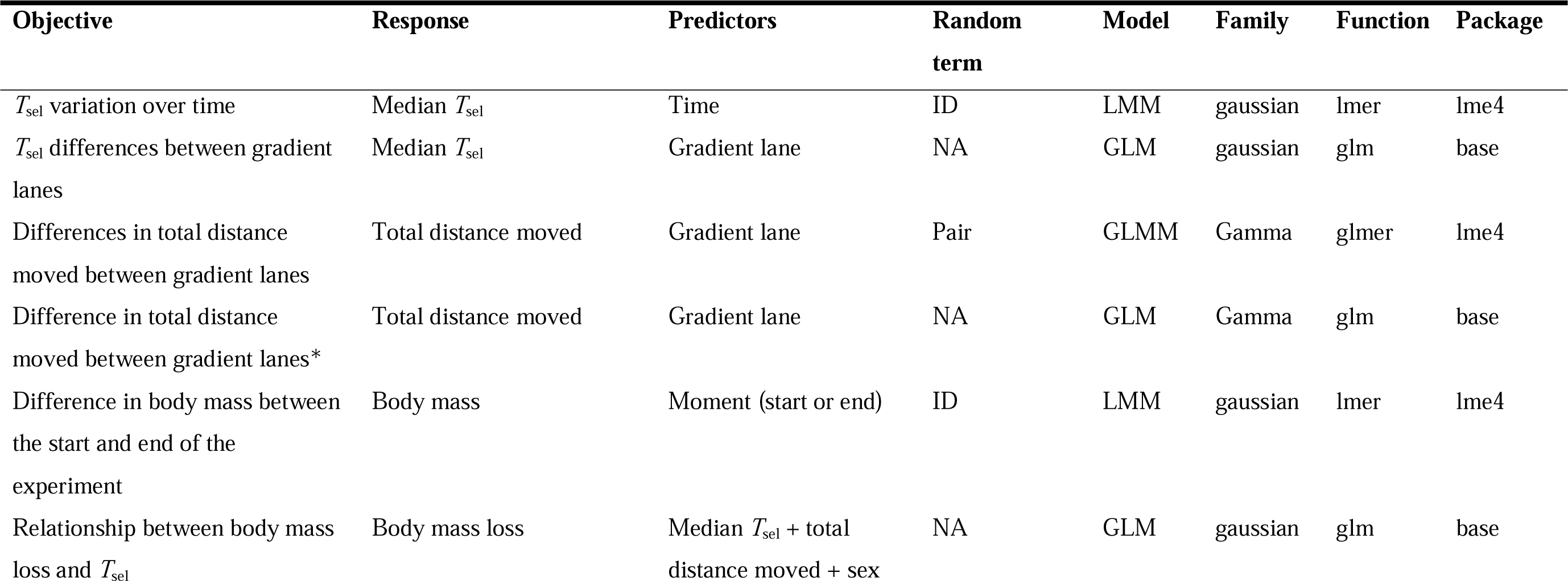
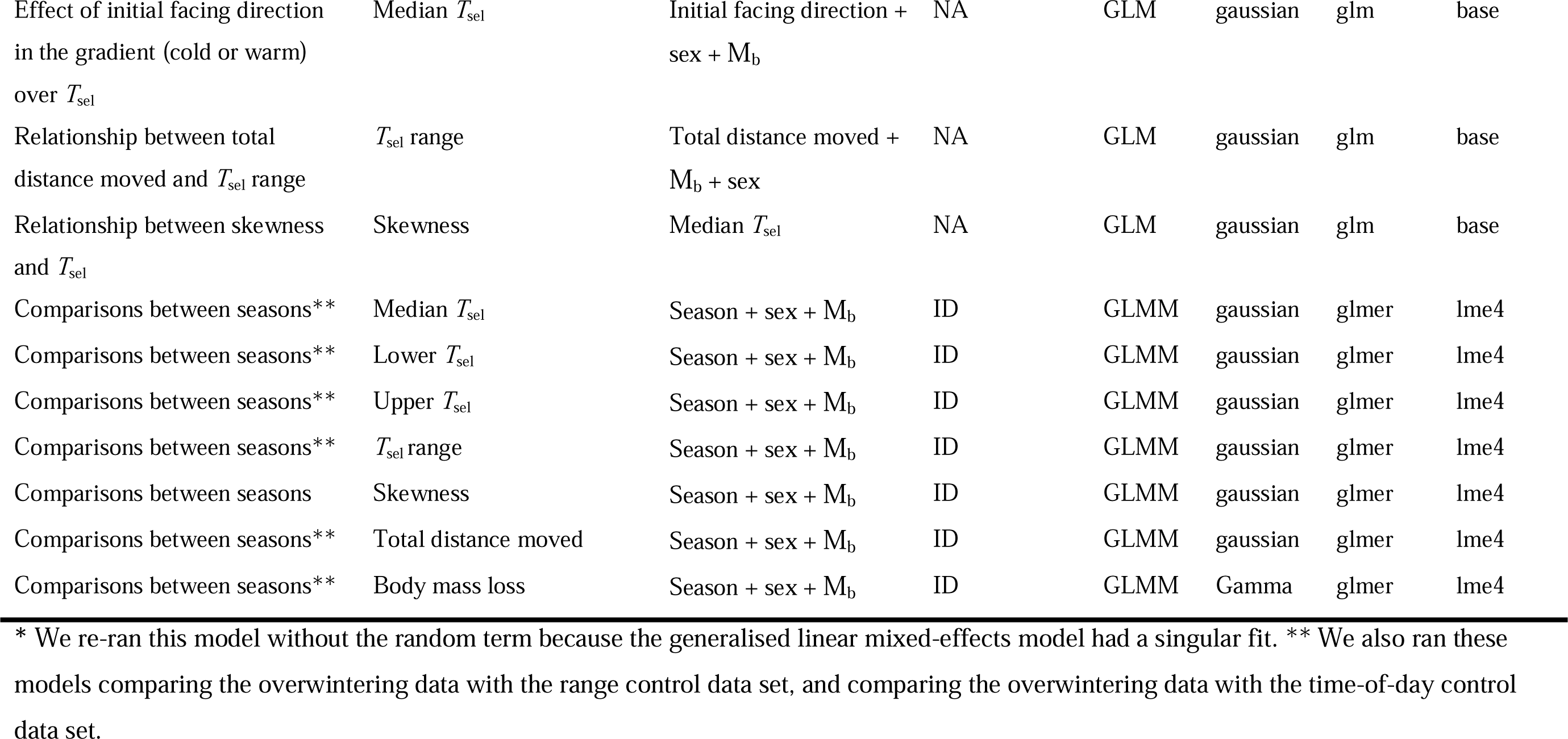
Statistical models tested in the current study. We provide information on the objective of each test, which variables constituted the models, the distribution family, as well as the functions and the R packages used to fit each model. Other than adjusting distribution families, we always used default parameters in the models. GLM = generalised linear model; LMM = linear mixed-effects model; GLMM = generalised linear mixed-effects models; *T*_sel_ = selected temperatures; M_b_ = body mass; ID = salamander identification.

**Table S2.**
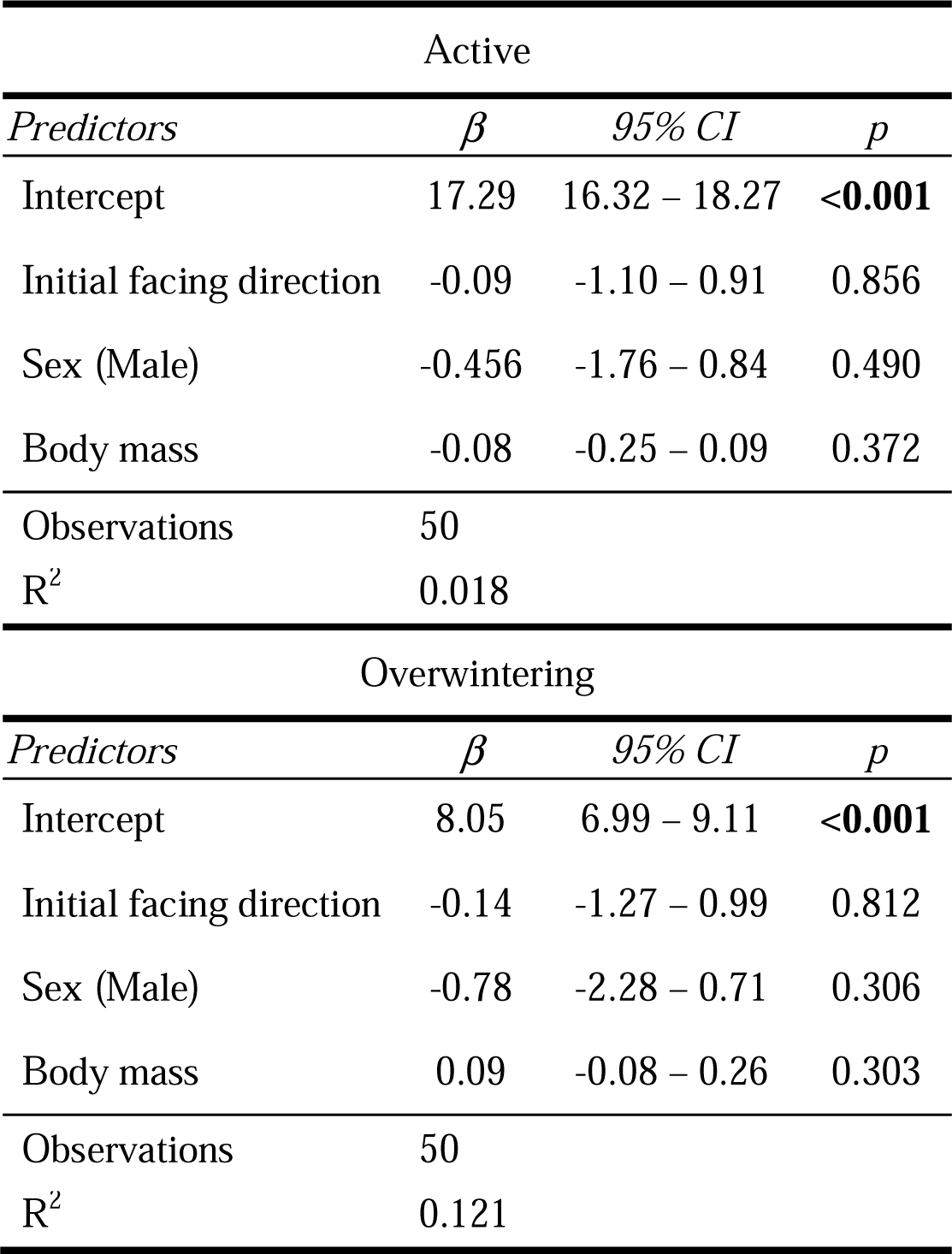
Parameter estimates (β), 95% confidence intervals (95% CI), and *p*-values for the models testing how the initial direction that *Ambystoma maculatum* faced in the thermal gradient affected their median selected temperature (*T*_sel_). The models had median *T*_sel_ as the response variable and facing direction, sex, and body mas as the predictors. Significant parameters are denoted in bold.

| Active |  |  |  |
| --- | --- | --- | --- |
| *Predictors* | $\beta$ | 95% CI | $p$ |
| Intercept | 17.29 | 16.32 – 18.27 | **<0.001** |
| Initial facing direction | -0.09 | -1.10 – 0.91 | 0.856 |
| Sex (Male) | -0.456 | -1.76 – 0.84 | 0.490 |
| Body mass | -0.08 | -0.25 – 0.09 | 0.372 |
| Observations | 50 |  |  |
| R<sup>2</sup> | 0.018 |  |  |
| Overwintering |  |  |  |
| *Predictors* | $\beta$ | 95% CI | $p$ |
| Intercept | 8.05 | 6.99 – 9.11 | **<0.001** |
| Initial facing direction | -0.14 | -1.27 – 0.99 | 0.812 |
| Sex (Male) | -0.78 | -2.28 – 0.71 | 0.306 |
| Body mass | 0.09 | -0.08 – 0.26 | 0.303 |
| Observations | 50 |  |  |
| R<sup>2</sup> | 0.121 |  |  |

**Table S3.** Parameters associated with temperature selection (*T*_sel_) and behavioural thermoregulation for a subset (N = 20) of *Ambystoma maculatum* studied in the overwintering seasons and those used as a range control. *p*-values in bold denote significant differences in a given trait between the active and overwintering season. SD = standard deviation; Min = minimum value; Max = maximum value; χ^2^= chi-square.

|  | Overwintering<br>(N=20) | Range control<br>(N=20) | Comparisons |
| --- | --- | --- | --- |
| **Median $T_{\text{sel}}$ (°C)** |  |  |  |
| Mean ± SD | 7.79 ± 2.17 | 7.30 ± 3.34 | $\chi^2_{(1)} = 0.41, p = 0.52$ |
| Median (Min, Max) | 7.64 (3.83, 13.20) | 7.97 (1.81, 13.70) |  |
| **Lower $T_{\text{sel}}$ bound (°C)** |  |  |  |
| Mean ± SD | 4.80 ± 2.82 | 5.42 ± 3.40 | $\chi^2_{(1)} = 0.64, p = 0.42$ |
| Median (Min, Max) | 4.11 (1.82, 13.20) | 4.29 (0.45, 13.40) |  |
| **Upper $T_{\text{sel}}$ bound (°C)** |  |  |  |
| Mean ± SD | 10.60 ± 1.80 | 8.85 ± 3.87 | $\chi^2_{(1)} = 3.24, p = 0.43$ |
| Median (Min, Max] | 10.90 (5.61, 13.40) | 9.23 (1.89, 14.60) |  |
| **$T_{\text{sel}}$ range (°C)** |  |  |  |
| Mean ± SD | 5.78 ± 2.70 | 3.43 ± 3.55 | $\chi^2_{(1)} = 7.63, p = \mathbf{0.03}$ |
| Median (Min, Max) | 6.85 (0.28, 8.59) | 1.80 (0, 9.90) |  |
| **Total distance moved (m)** |  |  |  |
| Mean ± SD | 92.70 ± 78.30 | 19.70 ± 37.00 | $\chi^2_{(1)} = 20.01, p < \mathbf{0.01}$ |
| Median (Min, Max) | 81.40 (0.20, 262) | 2.00 (0.01, 151) |  |
| **Body mass loss (%)** |  |  |  |
| Mean ± SD | 2.89 ± 2.35 | 2.91 ± 2.42 | $\chi^2_{(1)} = 0.0009, p = 0.97$ |
| Median (Min, Max) | 2.59 (0.11, 7.37) | 2.26 (0, 9.30) |  |

Note: we made comparisons between seasons using linear mixed effects models that had season, body mass, and sex as the predictor variables, as well as salamander ID as a random term. Given that neither body mass or sex mediated differences between seasons, we only report $\chi^2$ and $p$ -values for season in each model.

**Table S4.** Parameters associated with temperature selection (*T*_sel_) and behavioural thermoregulation for a subset (N = 20) of *Ambystoma maculatum* studied in the overwintering seasons and those used as a time-of-day control. *p*-values in bold denote significant differences in a given trait between the active and overwintering season. SD = standard deviation; Min = minimum value; Max = maximum value; χ^2^= chi-square.

|  | Overwintering<br>(N=20) | Time-of-day<br>control<br>(N=20) | Comparisons |
| --- | --- | --- | --- |
| **Median $T_{\text{sel}}$ (°C)** |  |  |  |
| Mean ± SD | 7.28 ± 2.17 | 6.78 ± 3.63 | $\chi^2_{(1)} = 0.29, p = 0.59$ |
| Median (Min, Max) | 7.12 (3.93, 13.10) | 7.88 (1.60, 11.70) |  |
| **Lower $T_{\text{sel}}$ bound (°C)** |  |  |  |
| Mean ± SD | 4.69 ± 2.40 | 5.28 ± 3.00 | $\chi^2_{(1)} = 0.47, p = 0.49$ |
| Median (Min, Max) | 4.33 (2.18, 13.0) | 4.79 (1.43, 11.0) |  |
| **Upper $T_{\text{sel}}$ bound (°C)** |  |  |  |
| Mean ± SD | 10.20 ± 2.12 | 7.74 ± 4.278 | $\chi^2_{(1)} = 4.40, p = 0.21$ |
| Median (Min, Max] | 10.60 (4.49, 13.10) | 9.47 (1.64, 12.90) |  |
| **$T_{\text{sel}}$ range (°C)** |  |  |  |
| Mean ± SD | 5.51 ± 2.81 | 2.45 ± 2.52 | $\chi^2_{(1)} = 14.17, p < \mathbf{0.01}$ |
| Median (Min, Max) | 6.14 (0.05, 9.16) | 1.20 (0.17, 7.10) |  |
| **Total distance moved (m)** |  |  |  |
| Mean ± SD | 71.20 ± 69.40 | 23.20 ± 31.60 | $\chi^2_{(1)} = 8.20, p = \mathbf{0.03}$ |
| Median (Min, Max) | 48.10 (0.25, 211) | 0.60 (0.03, 91.3) |  |
| **Body mass loss (%)** |  |  |  |
| Mean ± SD | 2.46 ± 1.57 | 3.39 ± 1.85 | $\chi^2_{(1)} = 3.07, p = 0.47$ |
| Median (Min, Max) | 2.07 (0.27, 5.80) | 3.24 (0.40, 7.33) |  |

Note: we made comparisons between seasons using linear mixed effects models that had season, body mass, and sex as the predictor variables, as well as salamander ID as a random term. Given that neither body mass or sex mediated differences between seasons, we only report $\chi^2$ and $p$ -values for season in each model.

### Supplementary figures

**Fig. S1.**
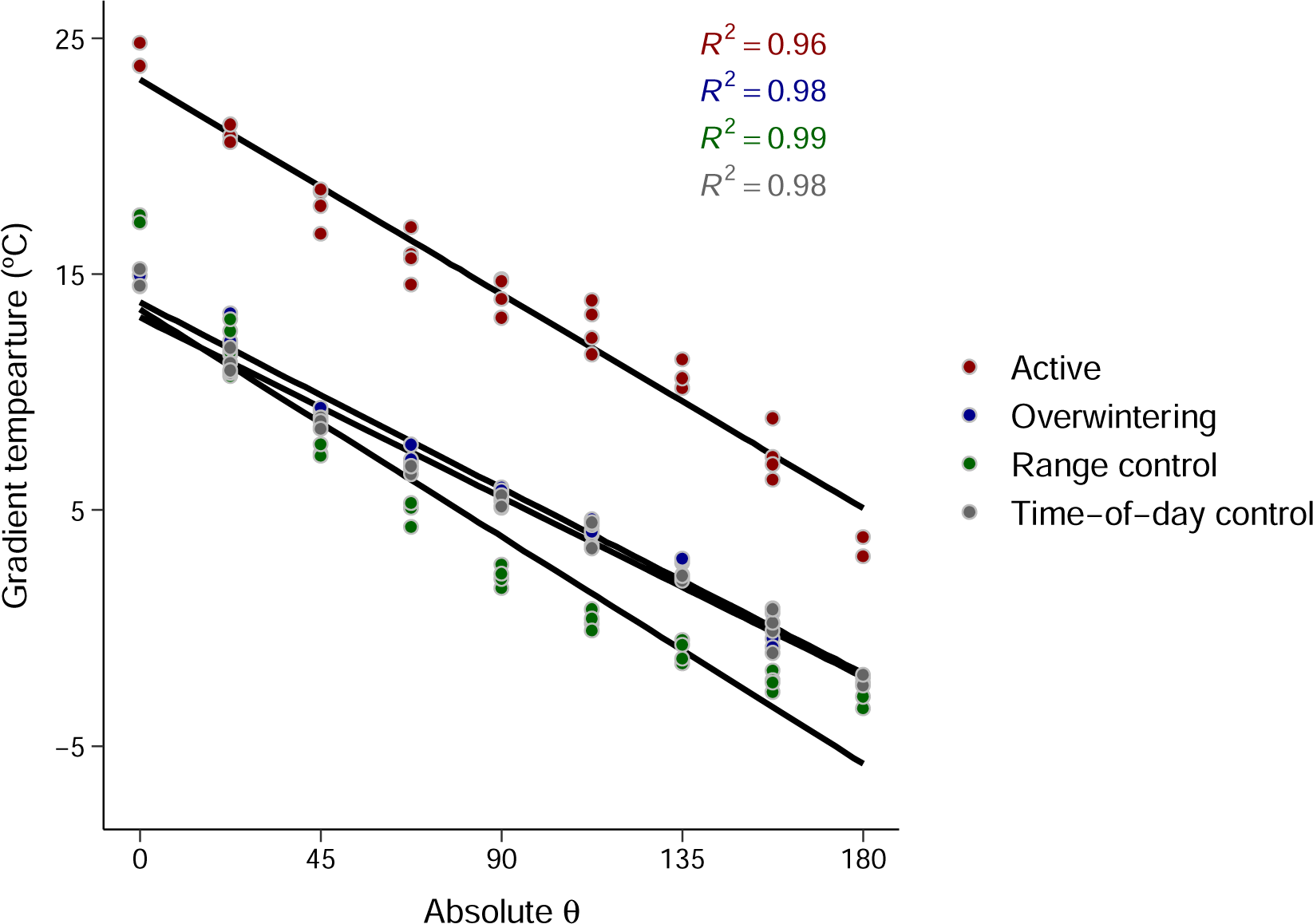
Relationship between gradient temperatures and polar coordinates (absolute θ). Seasons and their respective *R*^2^ values are colour-coded, with red indicating the active season, blue indicating the overwintering season, and green indicating the control. In the active season, the second-order regression equation describing the relationship between temperature and position in the gradient followed: y = 4_x_10^−5^x^2^ – 0.09x + 23.02. In the overwintering season, the second-order regression equation describing the relationship between temperature and position in the gradient followed: y = 8_x_10^−5^x^2^ – 0.10x + 14.24. In the range control experiments, the second-order regression equation describing the relationship between temperature and position in the gradient followed: y = 6 _x_10^−4^x^2^ – 0.21x + 16.65. In the time-of-day control experiments, the second-order regression equation describing the relationship between temperature and position in the gradient followed: y = 1_x_10^−4^x^2^ – 0.10x + 13.73. The

**Fig. S2.**
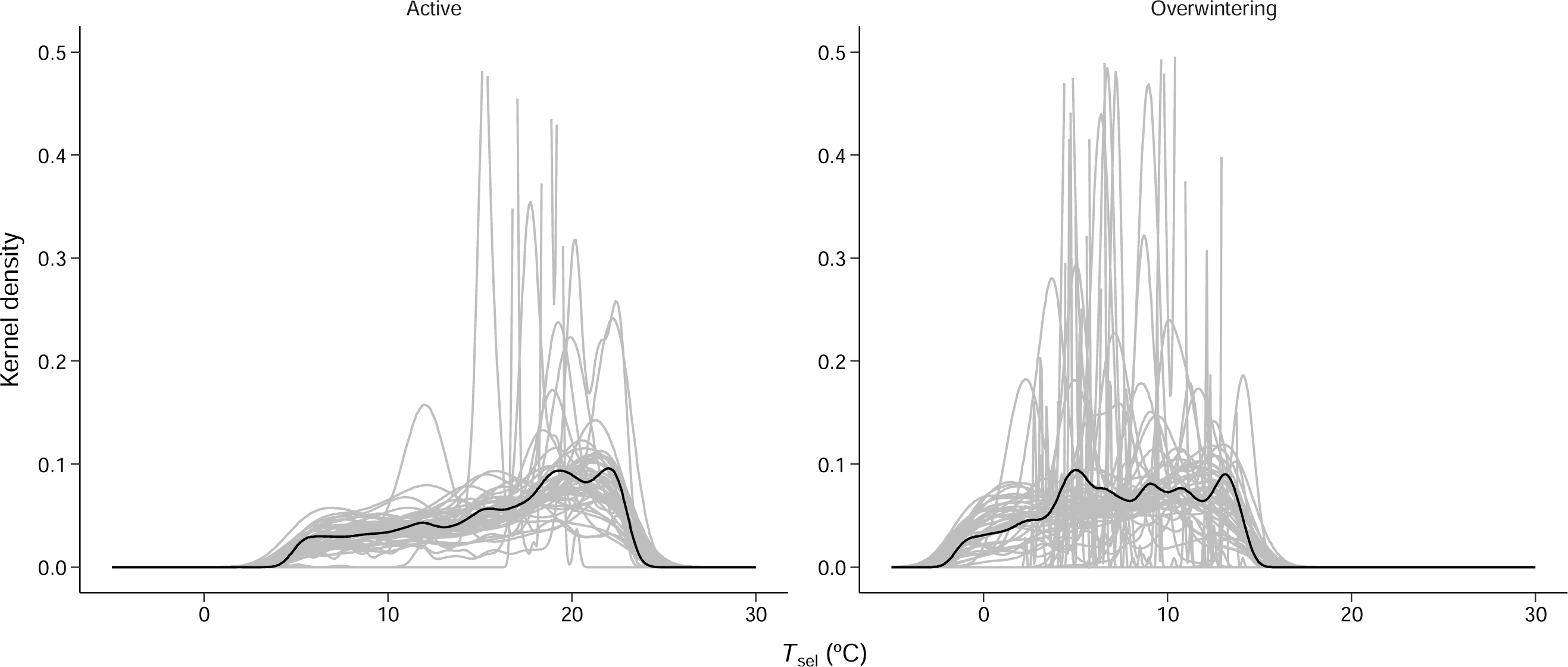
Kernel density estimations of median selected temperature (*T*_sel_) highlighting the negatively skewed and platykurtic nature of *T*_sel_ in *Ambystoma maculatum*. In both panels, the grey lines depict the *T*_sel_ distribution of each individual, whereas the black lines show the mean *T*_sel_ distribution considering all individuals tested within a season (N = 50 each season). Values above 0.5 on the y-axis pertain to salamanders that remained mostly stationary during the experiments.

